# Accelerating String Comparison in RLZ Compressed Sequences via LCE Jumps

**DOI:** 10.64898/2026.06.11.731742

**Authors:** Rahul Varki, Christina Boucher

## Abstract

Relative Lempel–Ziv (RLZ) is an effective compression method for large, repetitive collections; however, the fundamental primitives required to elevate it from a passive archival format to a tractable representation for compressed construction have yet to be fully established. In this paper, we introduce an algorithmic framework for structurally comparing and lexicographically sorting sequences of RLZ factors. We characterize when direct factor comparisons are necessary and when they can be bypassed using RLZ specific shortcuts. We further introduce a method for extending truncated factors into right-maximal matches, enabling the recovery of matching statistics from the RLZ parse. Experimentally, RLZ sorting achieved speedups of up to 3.93× over character-based sorting. Together, these results advance the use of the RLZ format as a foundation for compressed construction.

## 1 Introduction

As datasets continue to grow, data compression has become essential, motivating the need for compression methods whose output can support algorithms and queries directly without decompression. Relative Lempel–Ziv (RLZ) [18] is an algorithm that compresses a source text into a sequence of factors, each representing a right-maximal exact substring match in a smaller external reference text. It has been used successfully as an efficient compressor for biological sequences [18], textual documents [15, 21], and suffix arrays [8, 29, 30]. Beyond compression, RLZ representations have enabled random access [7, 18], self-indexes for pattern matching [9, 12, 27], and grammar compression [33]. Recently, the PIL-LAR model [5] was introduced to unify operations such as pattern matching and lexicographic sorting across standard, dynamic, and grammar-compressed strings by abstracting core primitives. Extending such an abstract framework to RLZ requires new techniques to structurally realign and compare factors between the source and reference without fully decompressing them.

To address this challenge, we introduce a method to structurally compare and lexicographically sort suffixes of a source text by operating directly on the RLZ parse using reference-based longest common extension (LCE_*R*_) queries. Given data structures supporting *O*(1)-time LCE_*R*_ queries, two RLZ factors can be compared lexicographically in constant time, substantially reducing the number of comparisons required relative to character-based methods. We identify when direct LCE_*R*_ queries are necessary and when comparisons can be resolved using factor ordering in the reference alone. We also show how non-maximal factors can be transformed into right-maximal factors to improve comparison efficiency. As a secondary theoretical consequence, this framework enables the direct construction of the *matching statistics* of the source text with respect to the reference from the RLZ parsing. Recently, Lipták et al. [23] showed that compressed matching statistics can be used directly for suffix sorting in highly similar string collections. In contrast, we begin with a RLZ representation and show that matching statistics can be recovered and exploited directly from RLZ factors, eliminating the need to store them explicitly while enabling compressed lexicographic comparison and sorting. Although motivated by suffix sorting, the framework applies more generally to lexicographic sorting of arbitrary sequences of RLZ factors. We implemented the method and evaluated it against standard character-based sorting on increasingly large collections of SARS-CoV-2 sequences [28]. The implementation is publicly available at https://github.com/rvarki/RLZ/.

## 2 Preliminaries

In this section, we establish the fundamental notation and concepts used throughout the paper.

### 2.1 Basic definitions

A string *T* is a finite sequence of symbols *T* = *T* [1..*n*] = *T* [1] · · ·*T* [*n*] over an alphabet *Σ* ={*c*_1_, … , *c*_*σ*_} whose symbols can be unambiguously ordered. We denote by *ε* the empty string, and the length of *T* as *T* . We denote by *T* [*i*..*j*] the substring *T* [*i*] · · · *T* [*j*] of | *T* | starting at position *i* and ending at position *j*, with *T* [*i*..*j*] = *ε* if *i > j*. For a string *T* and 1 ≤ *i* ≤ *n, T* [1..*i*] is called the *i*-th prefix of *T* , and *T* [*i*..*n*] is called the *i*-th suffix of *T* . A prefix *T* [1..*i*] is called a proper prefix if 1 ≤ *i < n*, while a suffix *T* [*i*..*n*] is called a proper suffix if 1 *< i* ≤ *n*. Let *A*_*B*_ denote the data structure *A* constructed over *B*.

### 2.2 Relative Lempel-Ziv

Relative Lempel-Ziv (RLZ) is an online dictionary-based compression scheme introduced by Kuruppu et al. [18]. At a high level, RLZ represents a source text *T* as a sequence of factors, each corresponding to a substring of a (smaller) reference text *R*. More specifically, RLZ parses the source text by greedily selecting, at each step, the longest prefix of the remaining unparsed text that matches a substring of the reference. Each factor is encoded as a (*p, ℓ*) pair, where *p* denotes the starting position of the match in the reference and *ℓ* the length of the match. Since the factors are chosen greedily, they are right-maximal with respect to the current position. We refer to these factors as *complete* factors. We define *incomplete* factors as segments that begin at an internal offset within complete factors. Formally, an incomplete factor is a proper suffix of a complete factor and can be represented as (*p* + *δ, ℓ* − *δ*), where 0 *< δ < ℓ*. We assume that all characters in the source appear in the reference (*Σ*_*T*_ ⊆ *Σ*_*R*_) and that identical substrings are parsed deterministically.

We illustrate the algorithm with the following example. Suppose that we apply RLZ to the source *T* [1..13] = AGCTCCAAGCTCC using the reference *R*[1..10] = AGCTCAAGC$. The resulting factorization is RLZ(*T* | *R*) = (1, 5), (5, 5), (4, 2), (3, 1). Together with the reference, the factorization is sufficient to reconstruct the source text.

### 2.3 Suffix Structures and Range Queries

#### SA and ISA

The suffix array (SA) of a text *T* [1..*n*] is an array of length *n* such that SA[*i*] stores the starting position of the *i*-th lexicographically smallest suffix of *T* [25]. The inverse suffix array (ISA) is its inverse permutation, where ISA[*i*] gives the lexicographic rank of the suffix starting at position *i* in *T* [25]. The data structures have the following relationship: ISA[SA[*i*]] = *i*.

#### LCP, RMQ, and LCE

The longest common prefix (LCP) array of a text *T* [1..*n*] is an array of length *n* in which LCP[*i*] denotes the length of the longest common prefix between the suffixes starting at SA[*i*] and SA[*i* − 1] for 1 *< i* ≤ *n* [25]. By convention, we set LCP[1] = 0 since there is no lexicographically smaller suffix. A range minimum query, RMQ_*A*_(*i, j*), returns the minimum value in the subarray *A*[*i*..*j*] for 1 ≤ *i* ≤ *j* ≤ |*A* |. There exist data structures that support *O*(1) RMQ queries [10]. The longest common extension query, LCE(*i, j*), returns the length of the longest common prefix between the suffixes *T* [*i*..*n*] and *T* [*j*..*n*] [19]. Let *r*_*i*_ = ISA[*i*] and *r*_*j*_ = ISA[*j*]. Assuming that *r*_*i*_ *< r*_*j*_, then: LCE(*i, j*) = RMQ_LCP_(*r*_*i*_ + 1, *r*_*j*_).

### 2.4 Matching Statistics

The matching statistics (MS) [3, 4] of *T* with respect to *R* is an array of length |*T*| , where MS[*i*] stores the length of the longest prefix of *T* [*i*..] that occurs in *R*. This array can be used to find maximal exact matches (MEMs) between *T* and *R*, which is critical in many tools [20, 31, 34, 2, 14, 1, 32, 35]. The RLZ parse can be viewed as a sparse representation of the MS array, sampled at the starting positions of each factor in the source text [23, 24].

## 3 Algorithmic Framework for RLZ Factor Comparison

This section presents our framework for sorting RLZ sequences directly. We first establish a theoretical foundation for comparing any two RLZ factors in *O*(1)-time using LCE_*R*_ queries supported by standard index structures over the reference. We then introduce optimization strategies that avoid explicit LCE_*R*_ evaluations by exploiting the lexicographic order of the reference text. Finally, we describe an efficient method for restoring right-maximality to incomplete RLZ factors. Pseudocode for the algorithms described in the main text is provided in the Appendix.

### 3.1 Constant-Time RLZ Factor Comparisons

A naive approach to lexicographically comparing two strings represented as RLZ sequences is to decompress them and perform a character-by-character scan in *O*(*n*)-time, where *n* is the length of the shorter string. However, because each RLZ factor corresponds to a substring in the reference, and any substring is simply a prefix of some suffix, we can bypass this character-level scan entirely. By precomputing the ISA_*R*_, LCP_*R*_, and 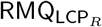 structures, the longest common prefix between any two factors can be determined in constant time via an LCE_*R*_ query. We note that while the use of LCE queries to evaluate compressed segments is known in internal pattern matching [17], it has not been formalized for RLZ factor sorting prior to this work. To the best of our knowledge, previous approaches to sorting RLZ sequences have fundamentally relied on string evaluation, either through character extraction or fingerprinting [9, 27, 11].

An LCE_*R*_ query allows the comparison to jump over matching segments, advancing to the next factor only when the current pair is exhausted. By maintaining relative offsets, transitions between factors take *O*(1)-time, bounding the total comparison to *O*(*r* + *q*)-time, where *r* and *q* are the numbers of factors in the two sequences. Because each LCE_*R*_ query advances the process by at least one factor, and *r* + *q* ≪ *n* when the reference captures the repetitive structure of the source, this factor-based sorting strategy significantly reduces the comparison overhead. Furthermore, when comparing complete factors of different lengths, the ordering is guaranteed to be resolved with a single LCE_*R*_ query, as established in Observation 1.

#### Observation 1.

*Let f*_1_ = (*p*_1_, *ℓ*_1_) *and f*_2_ = (*p*_2_, *ℓ*_2_) *be two complete RLZ factors of different lengths. Their lexicographic comparison requires at most one LCE*_*R*_ *query. Let k* = *LCE*_*R*_(*p*_1_, *p*_2_).

1. *If k <* min(*ℓ*_1_, *ℓ*_2_), *the order is determined by the characters R*[*p*_1_ + *k*] *and R*[*p*_2_ + *k*].
2. *If k* ≥ min(*ℓ*_1_, *ℓ*_2_) *and, w*.*l*.*o*.*g*., *ℓ*_1_ *< ℓ*_2_, *the order is determined by the first character of the factor following f*_1_ *in T and the character R*[*p*_2_ + *ℓ*_1_].

*Proof*. Case (1) is the standard definition of a mismatch in a string. For Case (2), assuming w.l.o.g that *ℓ*_1_ *< ℓ*_2_, then *f*_1_ is a prefix of *f*_2_. Since *f*_1_ is a complete factor produced by the greedy RLZ parser, it represents the longest possible match in the reference starting at its position. Therefore, the character immediately following *f*_1_ in the source text must differ from the character following *f*_1_ in the reference (which is the next character in *f*_2_), ensuring a mismatch.

### 3.2 Limitations of Ordering Strictly by the Reference

Although factors can be compared in *O*(1)-time using LCE_*R*_ queries, these queries incur a larger constant-factor overhead than direct character comparisons. This motivates the question of whether explicit LCE_*R*_ queries are necessary in this setting. Since each factor corresponds to a position in the reference, this raises the possibility of determining their lexicographic order directly from their ISA_*R*_ ranks, which represent the lexicographical ranks of the corresponding reference suffixes, thereby avoiding LCE_*R*_ queries altogether.

To test this hypothesis, we apply this approach to sort the suffixes of the source text prefixed by the complete RLZ factors from the example introduced at the end of Section 2.2, where RLZ(*T|R*) = (1, 5), (5, 5), (4, 2), (3, 1). Given that ISA_*R*_ = [4, 9, 7, 10, 6, 2, 3, 8, 5, 1], sorting the factors by the ISA_*R*_ ranks of their starting positions yields the ordering (1, 5), (5, 5), (3, 1), (4, 2). However, the correct lexicographic ordering is (1, 5), (3, 1), (5, 5), (4, 2).

We observe that the ordering is almost correct, except for the factor (3, 1). The issue is that this factor does not extend far enough into the reference for its starting position to uniquely determine its lexicographic order relative to the other factors. For instance, (3, 1) could alternatively be represented as (5, 1) or (9, 1), corresponding to the other occurrences of the factor in the reference. Since (3, 1) spans only one character, its ISA_*R*_ rank is determined solely by its right-context within the reference. In the source text, however, its true lexicographic rank is instead governed by the context introduced by the succeeding factor.

When comparing two RLZ factors, an LCE_*R*_ query identifies the exact point of divergence between their corresponding reference suffixes. If this LCE_*R*_ spans at least the length *ℓ* of either factor, that factor’s context in the reference becomes ambiguous. In this case, the distinguishing character lies outside the factor, so relying solely on its ISA_*R*_ rank may yield an ordering based on the reference suffix rather than its true continuation in the source text. Although the ISA_*R*_ alone cannot guarantee the absolute lexicographical order of all factors, it restricts their potential ranks to a localized interval within the SA_*R*_. In the next section, we exploit this approximate global ordering to establish a bound on the number of LCE_*R*_ comparisons needed to complete the sort.

### 3.3 Comparison Pruning via LCP-Induced SA Intervals

The lexicographic rank of a factor is determined by its starting position and length within the reference. As a factor extends deeper into the reference, its rank becomes increasingly coupled to the reference’s underlying suffix structure. This coupling allows factors with substantial right-context to be ordered directly using their ISA_*R*_ rank, avoiding the need for explicit LCE_*R*_ comparisons. We therefore classify factors as either *indicative* or *non-indicative*. We note that this bypassing of explicit evaluations is conceptually analogous to the lazy heuristics of CPS2 [6], whose underlying ideas have been further refined in modern LZ factorization [16], ensuring that LCE_*R*_ queries are executed only when necessary. Figure 1 highlights the indicative and non-indicative complete factors from the example introduced at the end of Section 2.2.

**Fig. 1.**
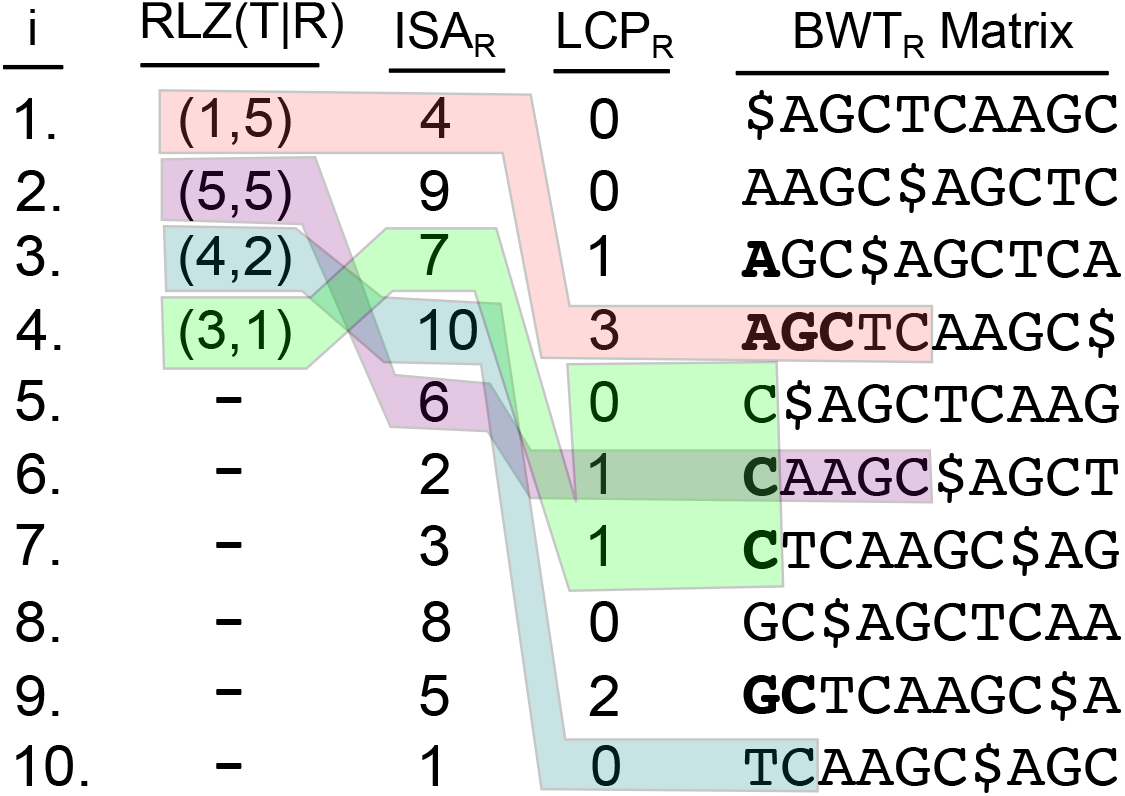
LCP_*R*_ intervals for indicative and non-indicative RLZ factors, using the example introduced at the end of Section 2.2. The columns show (from left to right) the index, RLZ factors, ISA_*R*_, and LCP_*R*_. The Burrows–Wheeler transform (BWT_*R*_) matrix is included for visualization, with characters contributing to the LCP_*R*_ matches shown in bold. Factors (1, 5), (5, 5), and (4, 2) are indicative, while (3, 1) is non-indicative. Indicative factors fully cover the bolded LCP_*R*_ value of their corresponding reference suffix, while non-indicative factors do not exceed it. The LCP_*R*_ intervals for (3, 1) and (5, 5) overlap, meaning LCE_*R*_ comparisons are required to determine their relative lexicographic order.

We define a factor as indicative if it corresponds to a unique match in the reference. Its ISA_*R*_ rank directly determines the position of the corresponding single-element interval in the SA_*R*_. As illustrated in Figure 1, the indicative complete factors in the example are (1, 5), (5, 5), and (4, 2). We can determine whether a factor is indicative in *O*(1)-time by checking if its length strictly exceeds the maximum LCP_*R*_ value its corresponding reference suffix shares with its immediate lexicographic neighbors; this follows from the fact that a suffix shares its maximal common prefix with its adjacent neighbors in the SA. Since these factors occupy single-element SA_*R*_ intervals, explicit LCE_*R*_ queries are only needed to break ties between indicative factors with the same ISA_*R*_ rank or when comparing against non-indicative factors whose SA_*R*_ intervals contain that rank.

We define a factor as non-indicative if it corresponds to multiple matches in the reference. Such a factor maps to a contiguous, multi-element interval in the SA_*R*_ containing at least two suffixes, meaning its ISA_*R*_ rank alone cannot resolve its exact position. In the worst case, this interval spans all SA_*R*_ positions whose corresponding reference suffixes begin with the factor’s first character. As illustrated in Figure 1, the only non-indicative complete factor in the example is (3, 1), which could alternatively be represented as (5, 1) or (9, 1). Although their distinct reference positions obscure this equivalence, the ISA_*R*_ resolves the ambiguity by mapping these identical matches to the contiguous SA_*R*_ interval spanning positions 5, 6, and 7. This contiguous grouping follows from the fundamental property that lexicographically identical prefixes correspond to adjacent suffixes in the SA. Thus, while a non-indicative factor’s ISA_*R*_ rank cannot pin-point its exact occurrence, it is guaranteed to fall within this broader interval. The exact boundaries of this interval are determined by the closest LCP_*R*_ values in either direction that are strictly less than the factor’s length. Given this ambiguity, explicit LCE_*R*_ queries are necessary for comparing factors with overlapping SA_*R*_ intervals, whereas factors with disjoint SA_*R*_ intervals can be ordered directly by their ISA_*R*_ ranks.

### 3.4 The Influence of Completeness on Factor Indicativity

The utility of a factor is closely related to whether it is complete or incomplete. An incomplete factor is truncated at its beginning and therefore captures only a suffix of a right-maximal match in the reference, providing less contextual information. In contrast, a complete factor preserves the entire right-maximal match. Consequently, complete factors are more likely to be indicative, often corresponding to a unique match in the reference, whereas incomplete factors are more likely to be non-indicative and correspond to multiple possible matches. We formalize this notion in Observation 2.

#### Observation 2.

*For a given RLZ parsing, complete factors are at least as indicative as any incomplete factors they cover*.

*Proof*. Let *f*_*c*_ be a complete factor and let *f*_*i*_ be an incomplete factor such that *f*_*i*_ is a proper suffix of *f*_*c*_. Consequently, the number of occurrences of *f*_*c*_ in the reference is at most the number of occurrences of *f*_*i*_ in the reference, since extending a match cannot increase the number of matching positions. This implies the following:

1. If *f*_*i*_ is indicative (i.e., appears exactly once in the reference), then *f*_*c*_, which occurs at a subset of those positions, must also appear exactly once in the reference and is therefore also indicative.
2. If *f*_*c*_ is non-indicative (i.e., appears multiple times in the reference), then *f*_*i*_ must appear at least as many times in the reference, and is therefore also non-indicative.

Suffixes of the source text prefixed by complete factors are the most efficient to order lexicographically, as they maximize the utility of the reference structure (Observation 2) and can themselves be efficiently ordered (Observation 1). Thus, the ordered set of source breakpoints with respect to the reference can be constructed efficiently. While the full impact of this structure on sorting efficiency remains open, it already accelerates comparisons involving incomplete factors. If the comparison advances both factors to their succeeding complete factors, the relative lexicographical order of the suffixes is determined directly by the breakpoint ordering.

### 3.5 Dynamic Factor Resynchronization

Incomplete factors are the most difficult to place lexicographically because they frequently map to non-indicative intervals. As discussed in Section 3.3, determining the rank of a non-indicative factor requires explicit LCE_*R*_ evaluation against a broader set of overlapping candidates. More critically, the fast-forwarding properties of Observation 1 do not hold for incomplete factors because they are not guaranteed to correspond to right-maximal matches in the reference. As residual substrings obtained by truncating complete matches, incomplete factors inherit the breakpoints of their parent factors rather than terminating at true reference mismatches. This introduces an artificial boundary at the end of the incomplete factor. Figure 2A illustrates an example of such an artificial breakpoint. While this results in only a single false breakpoint under standard RLZ parsing, it violates the right-maximal invariant and forces an unnecessary LCE_*R*_ query.

**Fig. 2.**
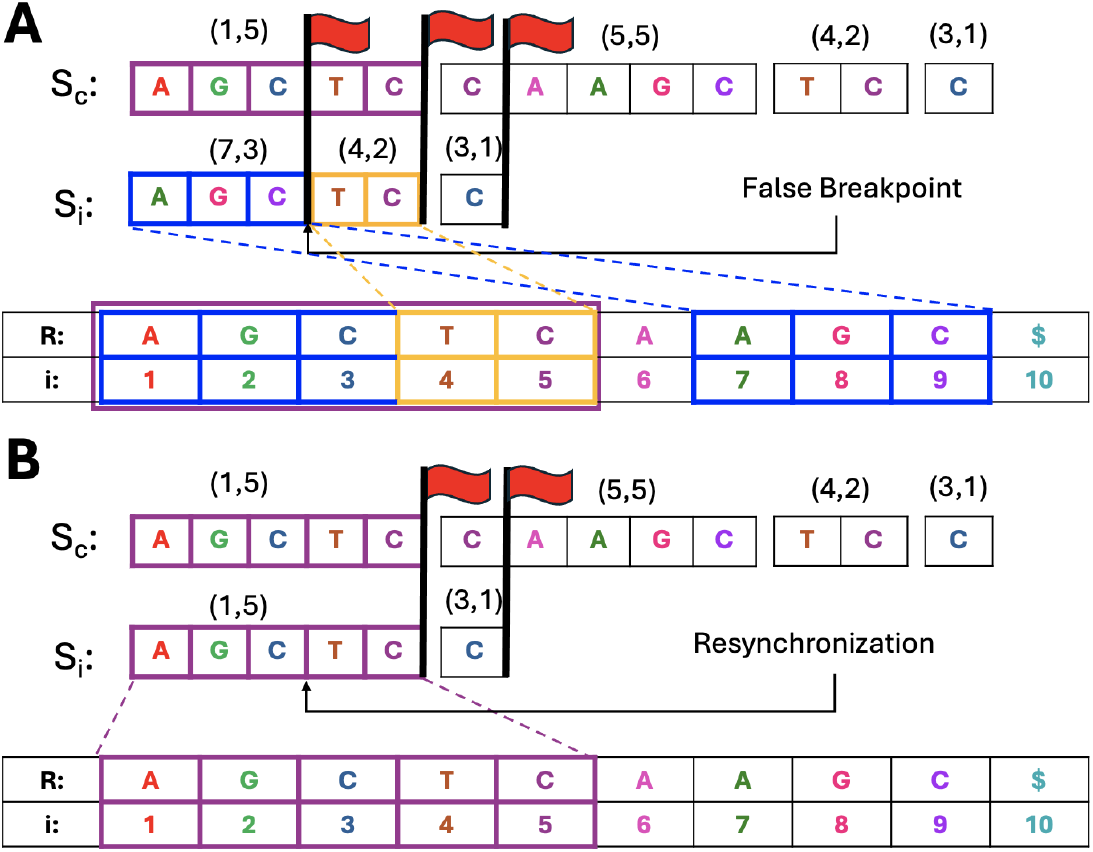
Synchronization issues arising from incomplete factors. In both subfigures, the suffix *S*_*c*_ = (1, 5), (5, 5), (4, 2), (3, 1) is being lexicographically compared against suffix *S*_*i*_ = (7, 3), (4, 2), (3, 1). *S*_*c*_ starts with a complete factor, whereas *S*_*i*_ starts with an incomplete factor. (A) Directly comparing the suffixes via their factors. This requires evaluating three breakpoints (denoted by red flags). The occurrences of the first two factors of *S*_*i*_ are also highlighted in the reference. Observe that one occurrence of each factor appears consecutively in the reference. (B) Application of the resynchronization method. By merging (7, 3) and (4, 2) into (1, 5), the artificial breakpoint (the first flag) is eliminated, restoring right-maximality with respect to the reference.

This computational overhead can be mitigated by dynamically merging adjacent factors. Instead of immediately halting at an artificial breakpoint, we search the incomplete factor’s lexicographic interval for a candidate position that can be extended by the successor. A successful extension indicates that the underlying reference match safely crosses the parsing boundary. We can determine how far an incomplete factor extends into its succeeding factor by finding the position in its SA_*R*_ interval that maximizes the LCE_*R*_ value between the residual suffix of the incomplete factor (i.e., excluding the prefix already matched by the factor) and the succeeding factor. By merging these factors to represent this extended match, we repair the boundary to guarantee that the resulting extended factor is right-maximal in the reference. This structural resynchronization eliminates false breakpoints, preventing compression artifacts from interrupting the comparison. Figure 2B illustrates this effect.

We note that while our resynchronization logic shares a common mathematical basis with the Longest Repeated Factor (LRF) heuristic used by Lipták et al. [22] to prune right-extensions during matching statistics construction, we use this property for a different structural purpose in RLZ. Instead of terminating an active search based on factor uniqueness, we exploit non-indicative suffix intervals to cross artificial parsing boundaries and dynamically resynchronize adjacent factors. As a further consequence, resynchronization enables direct construction of the matching statistics for the source text from the RLZ parsing. By restoring right-maximal alignment across all incomplete factors, we are implicitly computing the full matching statistics array, using complete factors as anchors. This is formalized in Theorem 1.

#### Theorem 1.

*Let T be a source text parsed into an RLZ sequence with respect to a reference R, where incomplete factors are implicitly covered by complete factors. Resynchronizing all incomplete factors restores their right-maximality, and together with complete factors yields the matching statistics of T with respect to R*.

*Proof*. By definition, the matching statistics (MS) of *T* with respect to *R* records, for each suffix of *T* , the length of its longest prefix in *R*. In the RLZ representation of *T* , each suffix is associated with either a complete or an incomplete factor. Complete factors are right-maximal in *R* by the greedy nature of the parse. Resynchronization restores right-maximality for incomplete factors by maximizing the LCE_*R*_ extension between residual suffixes in the SA_*R*_ interval of the incomplete factor and the succeeding complete factor. Hence, the procedure exactly computes the MS of *T* with respect to *R*.

## 4 Experiments

We evaluated our RLZ sorting methods against standard character-based sorting (hereafter denoted as *character sort*) for SA_*T*_ construction. The evaluation was performed on progressively larger, cumulative collections of SARS-CoV-2 sequences [28], containing up to 20,000 sequences. To ensure a fair algorithmic baseline, all implementations utilized `std::sort()`. We assessed four RLZ-based configurations: (i) *naive sort*, which performs factor comparisons using only LCE_*R*_ queries; (ii) *interval sort*, which leverages ISA_*R*_ ranks to bypass LCE_*R*_ evaluations whenever possible; (iii) *induced sort*, which proceeds in two passes: first sorting complete factors using ISA_*R*_ ranks, then sorting incomplete factors using the induced structure with ISA_*R*_ ranks where applicable, followed by a final merge; and (iv) *factor sort*, which sorts only the complete factors produced by RLZ using ISA_*R*_ ranks where applicable. We evaluated each configuration using both a single-sequence and a 1,000-sequence SARS-CoV-2 reference, neither of which was included in the datasets. We additionally evaluated all configurations with and without resynchronization. Naive, interval, and induced sorting decomposed complete factors into their incomplete factors to construct the full SA_*T*_ , whereas factor sort produced the subsampled SA_*T*_ corresponding to the suffixes prefixed by the complete RLZ factors. We excluded the preprocessing time required for factor decomposition, interval retrieval, and resynchronization from the reported sorting time, instead classifying these costs as preprocessing overhead since our primary focus was sorting performance. All experiments were run with Snakemake (v7.32.4) [26] on a server with 100 GB RAM and a AMD EPYC 75F3 32-core CPU clocked at 2.95 GHz. Preprocessing was parallelized using 16 threads, whereas sorting was performed on a single thread.

### 4.1 Empirical Efficiency of RLZ Sorting

We found that RLZ sorting can be significantly faster than character-based sorting, particularly with the shortcut-enabled configurations. As shown in Figure 3A, character sort required 21,517 seconds to sort all 20,000 SARS-CoV-2 sequences. Most RLZ sorting configurations improved upon this baseline, with naive sort taking 21,717 seconds, interval sort 20,096 seconds, induced sort 9,176 seconds, and factor sort 7,132 seconds. Note that these reported RLZ sorting times correspond to the configurations using a single-sequence reference.

**Fig. 3.**
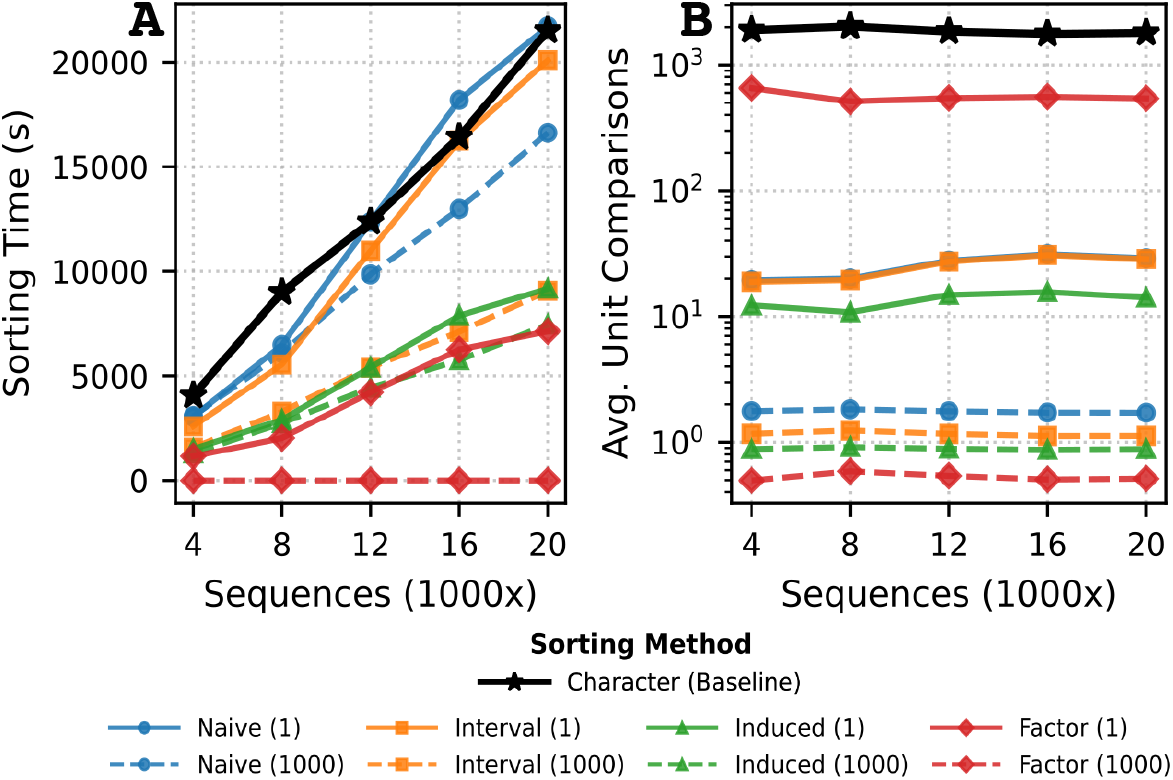
Character vs. RLZ sorting efficiency for SARS-CoV-2 sequences. (A) Sorting time (s) for constructing the SA_*T*_ . (B) Average unit comparisons per suffix comparison (character or factor). Brackets indicate the RLZ reference size.

Naive sort was slightly slower than character sort, while requiring significantly fewer factor comparisons. Figure 3B shows that all RLZ sorting configurations required fewer comparisons than character sort. This highlights the higher constant overhead of factor comparisons in practice, despite identical asymptotic time complexity. With interval sort, 47.9% of all suffix comparisons were resolved directly using SA_*R*_ intervals. For induced sort, 35.2% of the comparisons that required direct LCE_*R*_ evaluation were resolved using the sorted complete factor backbone. Factor sort, while the fastest configuration, had the highest average number of factor comparisons per suffix comparison (542) among all RLZ sorting configurations. This suggests that mismatches typically occur far apart in this dataset, particularly for suffixes prefixed by complete factors.

Sorting time for all RLZ configurations improved with the 1,000-sequence reference. As shown in Figure 3A, naive sort decreased to 16,625 seconds (23.4% decrease), interval sort to 9,067 seconds (54.9% decrease), induced sort to 7,457 seconds (18.7% decrease), and factor sort to 1 second (99.99% decrease) when sorting all 20,000 SARS-CoV-2 sequences. Figure 3B shows a similar decrease in the average number of factor comparisons, which drops from a range of 14–542 to 0.6–1.8 comparisons across configurations. As expected, these improvements are due to the significantly larger average factor length obtained with the 1,000-sequence reference, with the average factor length increasing from 34 ± 256 characters to 1,809 ± 3,007 characters. While using a larger reference is seemingly a natural choice given these results, the primary downside is increased memory usage.

### 4.2 Resynchronization Reduces RLZ Sorting Time

We found that resynchronizing incomplete factors into right-maximal factors prior to sorting improved overall sorting time for each applicable RLZ sorting configuration, particularly when using a larger reference. For the parsing generated with a single-sequence reference, resynchronization had a negligible effect on both sorting time and the average number of factor comparisons per suffix comparison (Figure 4A–B). Sorting time improved by only a few hundred seconds, while the average number of factor comparisons decreased by less than 0.01 across all subsets and configurations. In contrast, for the 1,000-sequence reference, resynchronization noticeably decreased sorting time and the average number of comparisons (Figure 4C–D). When sorting the full 20,000 SARS-CoV-2 sequences, naive sort decreased to 13,963 seconds (16.0% decrease), interval sort to 6,999 seconds (22.8% decrease), induced sort to 5,465 seconds (26.7% decrease), and factor sort to 1 second (0% decrease), compared to configurations without resynchronization enabled. Additionally, the average number of factor comparisons decreased by 0.22-0.30 factors per suffix comparison.

**Fig. 4.**
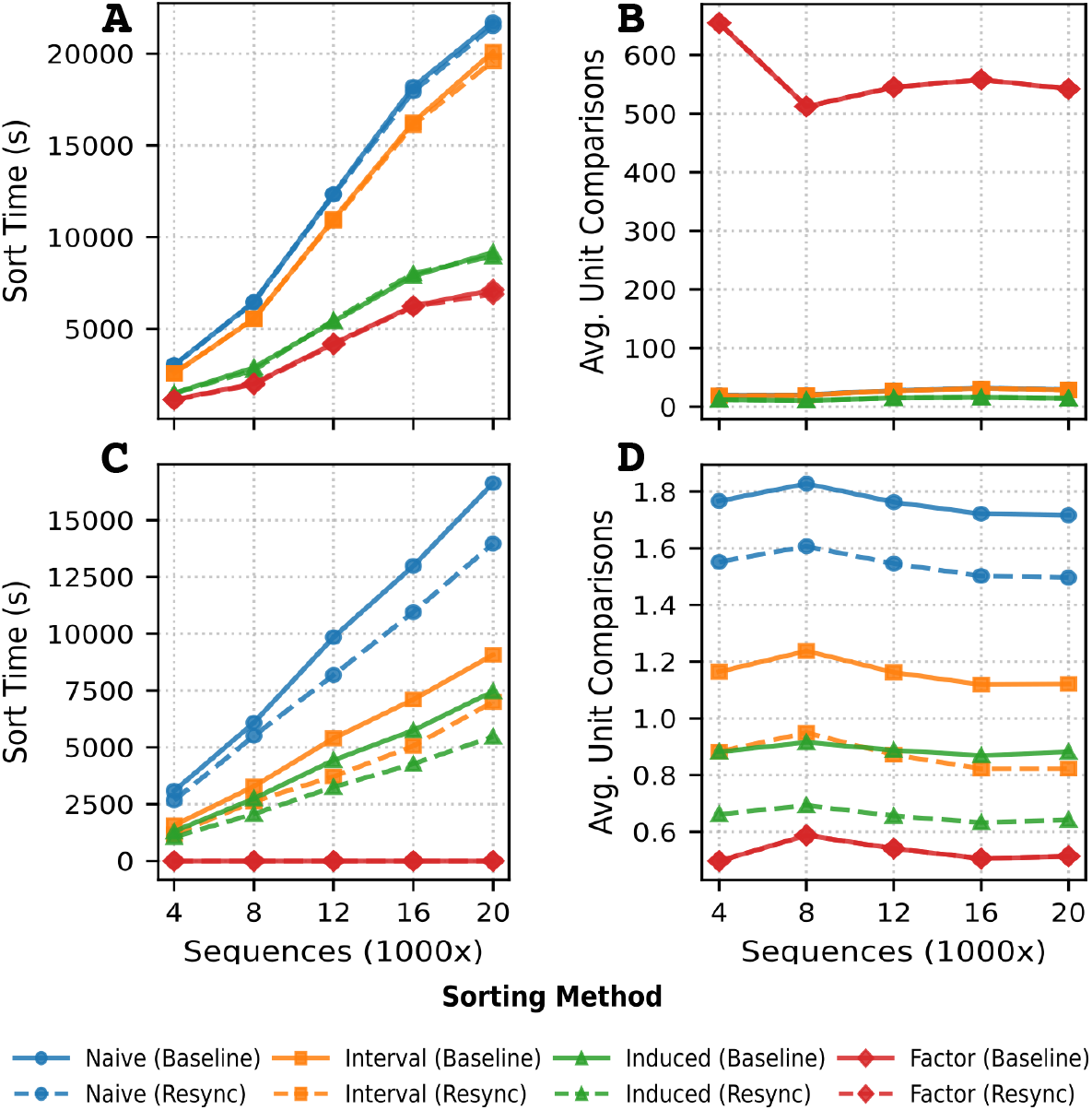
RLZ sorting efficiency with resynchronization for SARS-CoV-2 sequences. (A) Sorting time (s) for constructing the SA_*T*_ with the single-sequence reference. (B) Average factor comparisons per suffix comparison with the single-sequence reference. (C) Sorting time (s) with the 1,000-sequence reference. (D) Average factor comparisons per suffix comparison with the 1,000-sequence reference.

The larger reference provided significantly more extension points for incomplete factors, which explains why the resynchronization impact differed. Specifically, the resynchronization rate increased from 1.5% for the single-sequence reference parsing to 37.1% for the 1,000-sequence reference parsing. As a downstream effect of resynchronization, more factors became indicative and were therefore able to use the interval comparison shortcut more often. In general, while larger references often result in lower initial utilization, as a larger fraction of the reference remains unused, they often enable more effective resynchronization.

## 5 Conclusion

This paper introduces a novel algorithm that enables direct lexicographical comparison of RLZ factors and their compressed sequences. By constructing a small set of auxiliary structures over the reference text, our method resolves comparisons efficiently using LCE_*R*_ queries, while strategically leveraging the underlying reference structure to bypass explicit evaluations whenever possible. To further optimize performance, we presented a resynchronization algorithm that restores right-maximality to incomplete factors, thereby minimizing redundant LCE_*R*_ queries. As a secondary theoretical contribution, we also show how the matching statistics can be computed from the RLZ parse. Our experimental results demonstrate that RLZ sorting can outperform conventional character-based sorting, achieving speedups of up to 3.93 ×.

Several open questions warrant future investigation. First, can we directly construct the SA_*T*_ from the subset of entries corresponding strictly to complete factors? Furthermore, does a structure analogous to the *ϕ*-function for the r-index [13] exist for RLZ representations? Second, the optimal selection of a reference text for RLZ remains an open challenge, particularly regarding the trade-offs between factor length, reference size, and retained redundancy. As lexicographic comparison is a fundamental primitive in many algorithms, this work represents a step towards establishing RLZ not merely as a compression method, but as a practical, compute-ready representation.

## Acknowledgments

R.V. and C.B. were supported by “Fully Realizing Pangenomics Alignment” (NIH Award No. 1R56HG013865; Contact PI: Christina Boucher) and “Personal and Panel References for Improved Alignment” (NIH Award No. 2R01HG011392; formerly 5R01HG011392; Contact PI: Ben Langmead).

## 6 Appendix

The following pseudocode presents the algorithms referenced in the main text. Algorithm 1 describes the procedure for lexicographically comparing two RLZ sequences with LCE_*R*_ queries. Algorithm 2 determines whether a given factor is indicative of a unique match in the reference. Algorithm 3 computes the SA_*R*_ range corresponding to all occurrences of a non-indicative factor in the reference. Algorithm 4 details how to extend incomplete factors into right-maximal matches in the reference.

### Algorithm 1 Lex. comparison of RLZ sequences

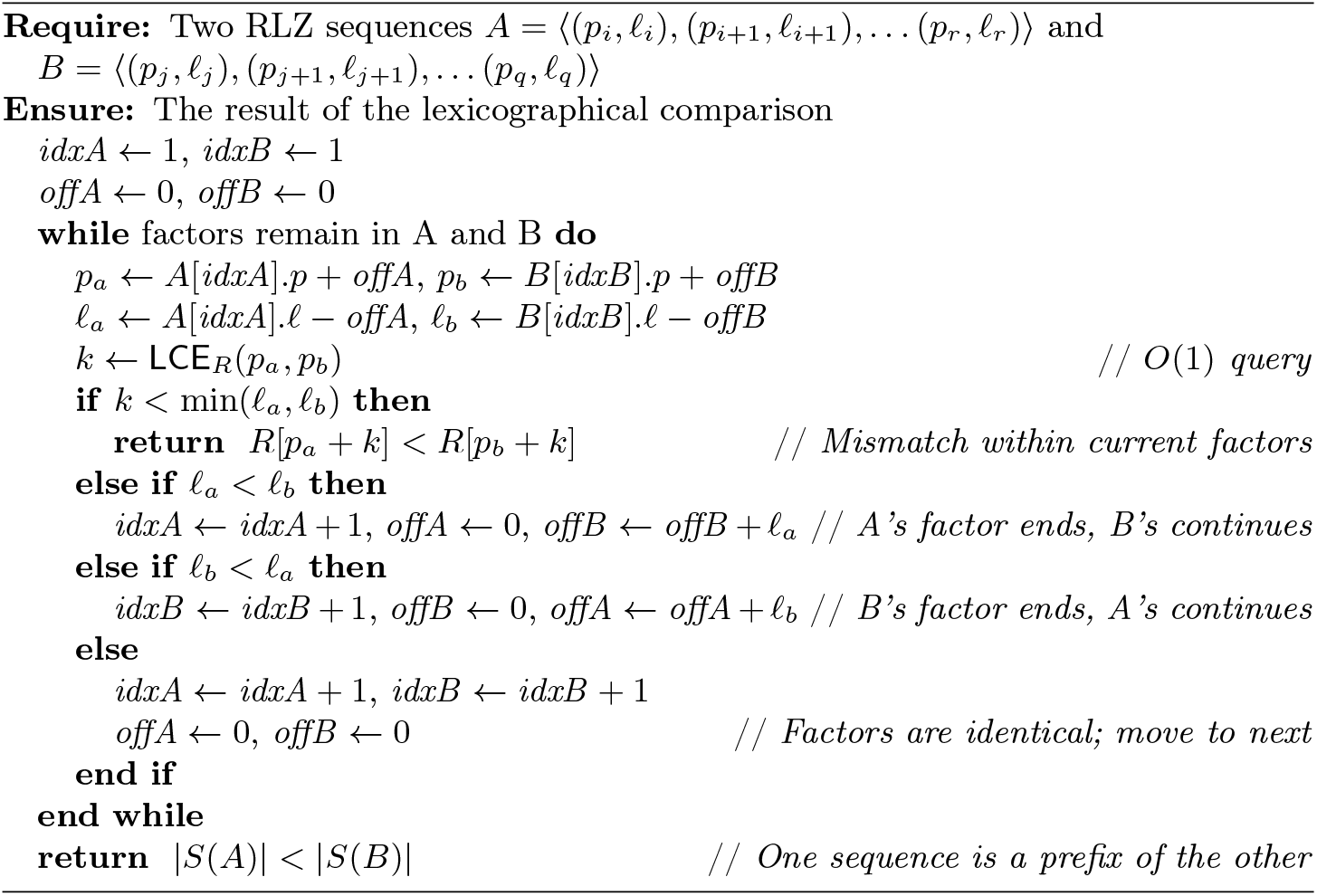

### Algorithm 2 Determine if factor is indicative

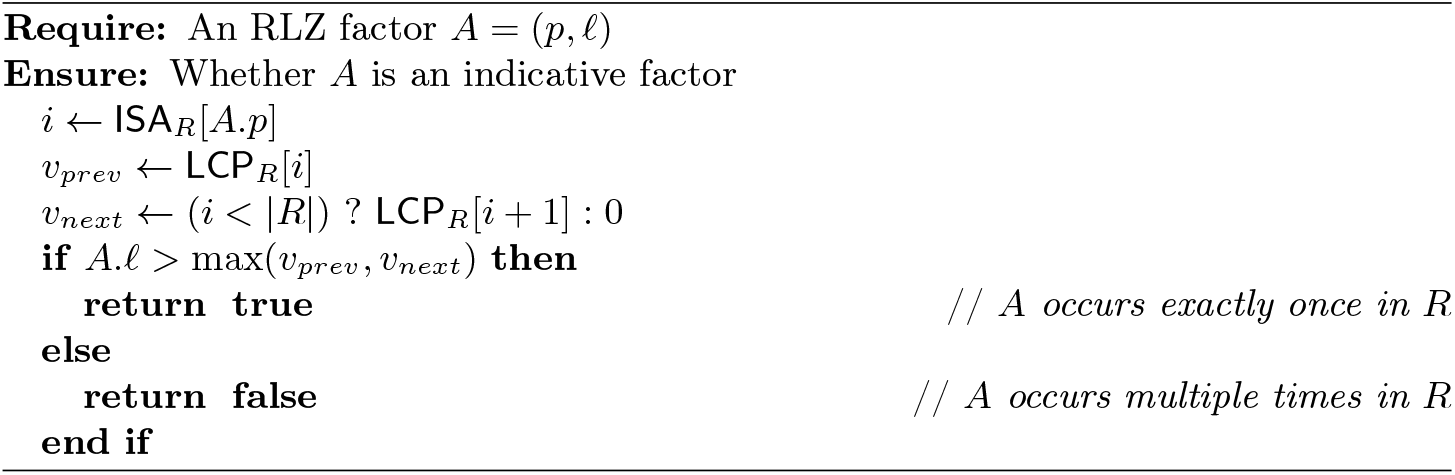

### Algorithm 3 Find SA_*R*_ interval of incomplete factor

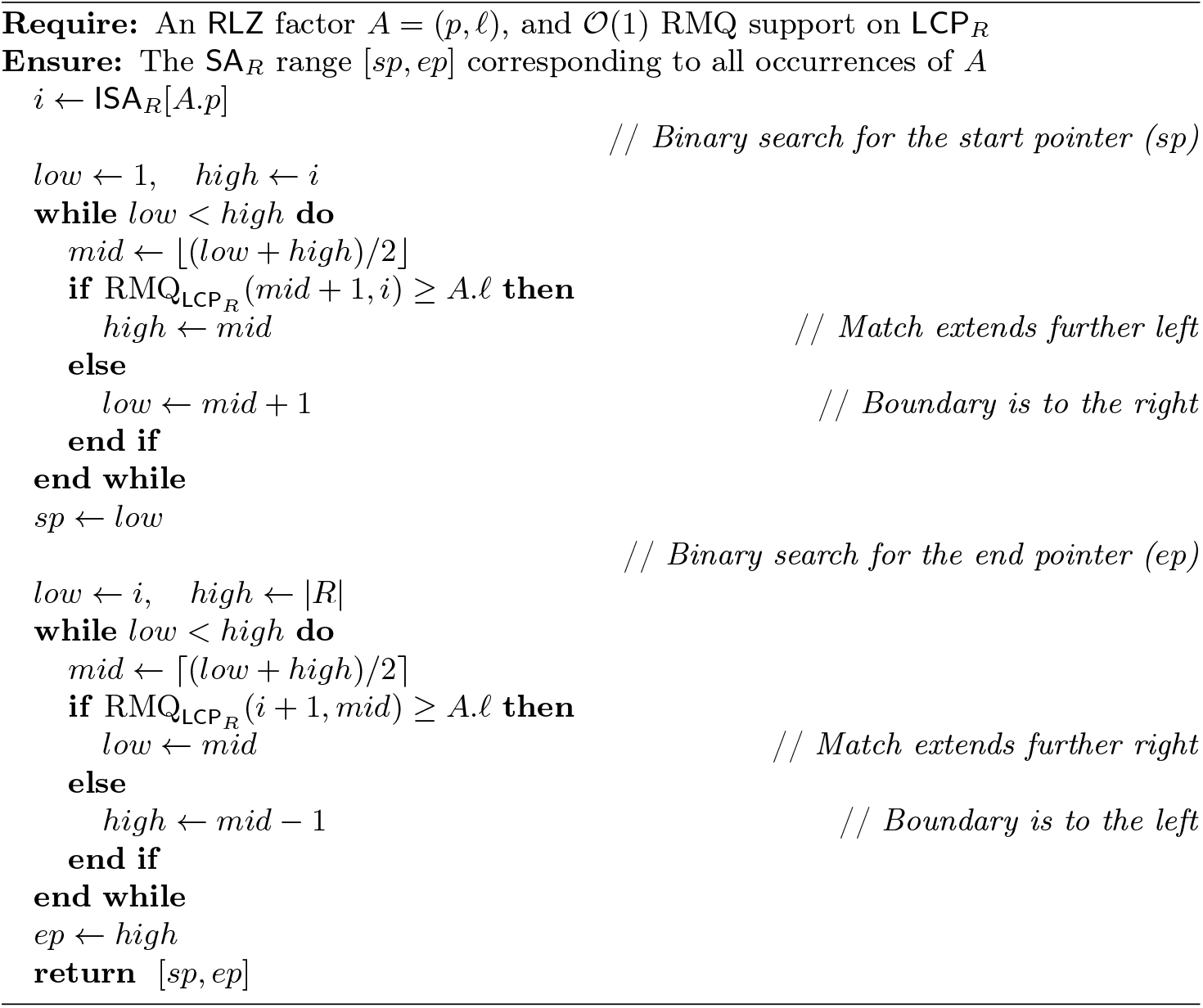

### Algorithm 4 Resynchronize outdated breakpoints

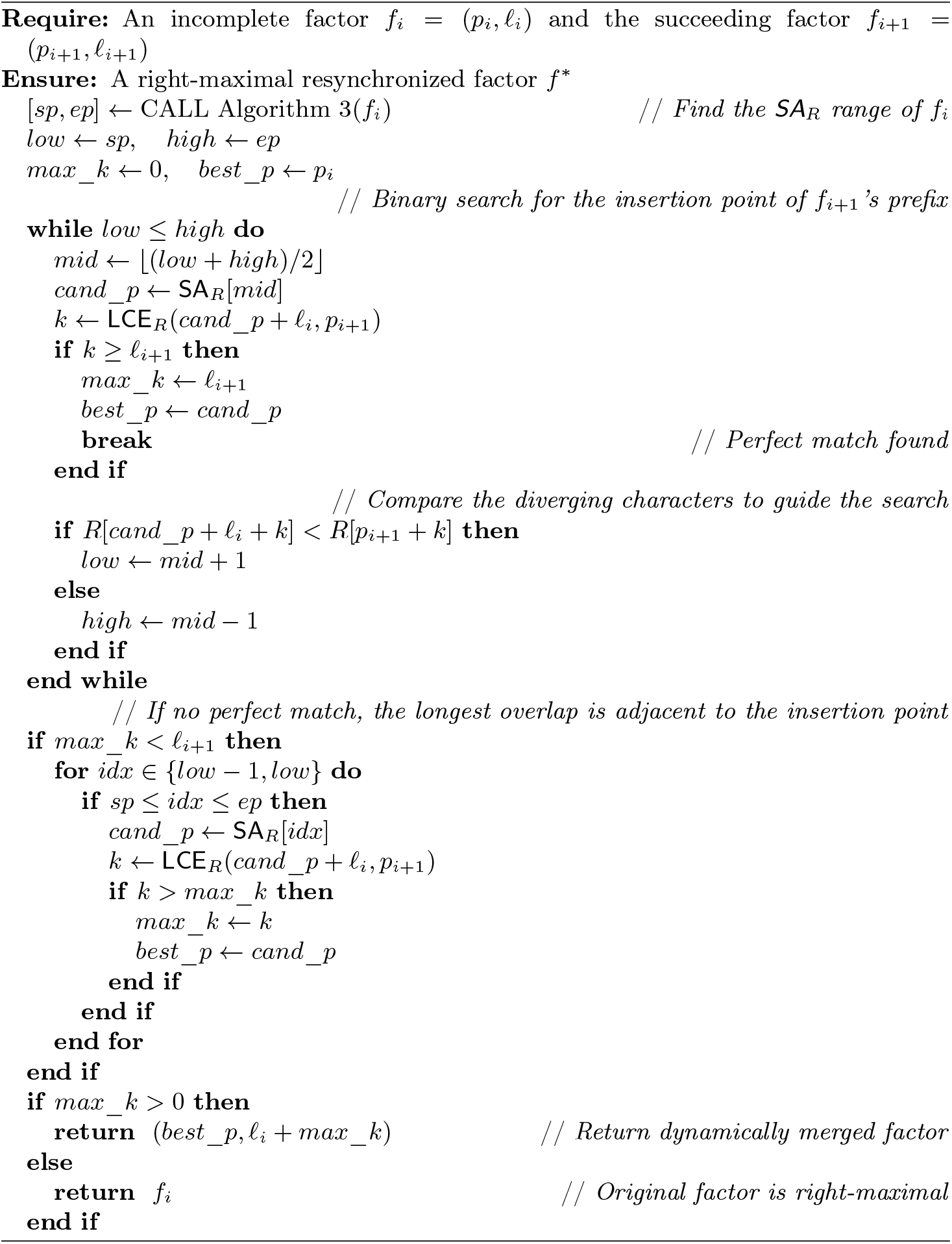

